# Probing Affinity, Avidity, Anti-Cooperativity, and Competition in Antibody and Receptor Binding to the SARS-CoV-2 Spike by Single Particle Mass Analyses

**DOI:** 10.1101/2021.06.18.448939

**Authors:** Victor Yin, Szu-Hsueh Lai, Tom G. Caniels, Philip J.M. Brouwer, Mitch Brinkkemper, Yoann Aldon, Hejun Liu, Meng Yuan, Ian A. Wilson, Rogier W. Sanders, Marit J. van Gils, Albert J.R. Heck

## Abstract

Determining how antibodies interact with the spike (S) protein of the SARS-CoV-2 virus is critical for combating COVID-19. Structural studies typically employ simplified, truncated constructs that may not fully recapitulate the behaviour of the original complexes. Here, we combine two single particle mass analysis techniques (mass photometry and charge-detection mass spectrometry) to enable measurement of full IgG binding to the trimeric SARS-CoV-2 S ectodomain. Our experiments reveal that antibodies targeting the S-trimer typically prefer stoichiometries lower than the symmetry-predicted 3:1 binding. We determine that this behaviour arises from the interplay of steric clashes and avidity effects that are not reflected in common antibody constructs (i.e. Fabs). Surprisingly, these sub-stoichiometric complexes are fully effective at blocking ACE2 binding despite containing free receptor binding sites. Our results highlight the importance of studying antibody/antigen interactions using complete, multimeric constructs and showcase the utility of single particle mass analyses in unraveling these complex interactions.

## Introduction

The emergence of the SARS-CoV-2 coronavirus and subsequent onset of the coronavirus disease 2019 (COVID-19) pandemic has necessitated the rapid development of vaccines and other treatments.^1–3^ The primary focus of these countermeasures is the SARS-CoV-2 spike (S) protein present on the viral surface, which is responsible for initiating host infection *via* complexation to the human ACE2 receptor and subsequent fusion of the viral and host cell membranes.^4^ The majority of vaccines developed against SARS-CoV-2 use the S protein (e.g. genetically encoded via either mRNA/DNA cargo^5–7^ or displayed on a nanoparticle surface^8^) to elicit an immune response. Understanding how exactly antibodies (Abs) interact with the SARS-CoV-2 S protein is a crucial component for both continuing vaccine development as well as the rational design of target biotherapeutics (e.g. monoclonal Abs).^9,10^

Like the spike proteins of many other viruses, the SARS-CoV-2 S protein is present in a trimeric, membrane-embedded state.^11^ Effective neutralizing Abs for SARS-CoV-2 often target the receptor binding domain (RBD) of the S protein.^12–15^ As the RBD is the site of initial ACE2 receptor binding, these Abs are thought to achieve neutralization largely by sterically preventing interactions between the S protein and host receptor.^16^ Due to its trimeric nature, each individual spike contains three copies of the RBD.

Given the central role of Ab binding for the successful neutralization of antigens, a seemingly simple question is: how many copies of an Ab can bind to one spike? And relatedly, how many Ab copies need to bind to induce neutralization? Since each S-trimer contains three identical copies of the S protomer, one may expect that Abs bind the S-trimer with a 3:1 stoichiometry. However, this prediction may be somewhat naïve, and the true Ab binding stoichiometry will be complicated by several factors. Firstly, the RBD is dynamic and can occupy either an “up” or “down” state, defined by its position relative to the remainder of the complex.^11^ Only the up RBD state is capable of binding the ACE2 receptor.^17^ As each RBD is related in the S-trimer by 3-fold symmetry, there exists a total of 4 possible conformational states of the RBDs in the S-trimer (with up : down ratios of 0:3, 1:2, 2:1, and 3:0). Certain Abs against the RBD may only recognize one of the two states, which can interconvert.^12,18,19^ Therefore, any RBD-targeting Ab could conceivably bind a particular S-trimer with any stoichiometry between 0 and 3, depending on the exact conformational status of the complex. Secondly, full Abs (IgGs) possess two equivalent Fab arms, of which one or both may be involved in binding (i.e. avidity). Avidity effects are well-known to play key roles in the potency of neutralizing Abs and could manifest as an apparent decrease in binding stoichiometry.^20,21^ Thirdly, anti-cooperative binding effects arising from steric conflicts between multiple binding Abs may also play a role, hampering the amount of concurrent binding allowed.

Considering the known impacts these various effects can have on Ab efficacy, the stoichiometries of Ab binding to the SARS-CoV-2 S protein is surprisingly poorly characterized. This is likely due in part to the lack of biochemical and biophysical methods to effectively probe such heterogeneous interactions effectively and efficiently. For example, surface plasmon resonance (SPR) and biolayer interferometry (BLI) are highly effective at rapidly quantifying antigen binding, but provide only an ensemble-averaged overview and yield limited structural information.^22–24^ Single particle electron microscopy (EM) can often provide near-atomic details of protein structure and protein-protein interactions (allowing direct mapping of Ab epitopes on the full SARS-CoV-2 S ectodomain^11,25–27^), but due to the extended flexibility of full-length IgGs is typically (with some exceptions^28^) only able to visualize binding of antibody fragments (i.e. truncated Fab domains), and thus may not directly capture any effects of avidity or steric interactions that would occur in the full IgG. Nuclear magnetic resonance spectroscopy and X-ray crystallography can yield atomic protein structures, but due to limitations with size and conformational/glycosylation-induced heterogeneity, respectively, have been largely restrained to studies on truncated single RBD constructs, and thus remain relatively blind to both the up : down dynamics of the full trimer as well as potential avidity effects.^29,30^

Native mass spectrometry (MS) is an analytical technique capable of measuring the mass of proteins and protein complexes.^31^ As any binding event leads to a corresponding increase in mass, native MS offers a convenient readout of ligand binding and can readily distinguish different binding stoichiometries and different ligands by their unique masses. In the context of monitoring interactions to the SARS-CoV-2 S protein, the feasibility of these experiments is greatly hindered by the extreme heterogeneity caused by the high degree of glycosylation present on the S protein (the so-called glycan shield).^32–34^ This heterogeneity leads to a normally untenable degree of spectral complexity that obfuscates the charge state assignments required for correct mass determination.^35^ While some success has been reported in the conventional native MS analysis of SARS-CoV-2 S and other viral spike proteins (e.g. by metabolic glycan engineering^36^ or limited charge reduction^37^ of truncated constructs), these modified constructs may not exhibit the same binding behaviour as the real viral spike protein, given the known importance of glycan structure in these interactions.^32^

Here, we report the application of two single particle approaches for mass analysis, mass photometry^38^ and charge-detection native mass spectrometry^39,40^, to circumvent the need of conventional charge assignment and allow successful measurement of the full SARS-CoV-2 S-trimer ectodomain, as well as the binding stoichiometries to full-length neutralizing IgGs. Our measurements reveal that IgG binding to the SARS-CoV-2 S-trimer can exhibit a diversity of binding behaviours that is not captured when studying the truncated Fabs or RBD constructs alone. We also demonstrate that these techniques can be used to monitor binding of the ACE2 receptor, as well as the S proteins from other variants of concern of the SARS-CoV-2 virus. These ultrasensitive single particle approaches (requiring only ∼femtomoles of sample) thus offer a powerful addition to the toolkit of contemporary biophysical tools by providing a “one-shot” method for determining Ab affinity, anti-cooperativity, and avidity simultaneously. Our findings highlight the biophysical complexity of the multimeric interactions that occur between Abs and the SARS-CoV-2 S protein.

## Results

### Single Particle Mass Analysis of the SARS-CoV-2 S-trimer

Originally introduced as interferometric scattering mass spectrometry (iSCAMS)^38^, mass photometry (MP) is a light scattering-based, label-free, mass analysis technique that determines the mass of a single particle from its scattering intensity.^41^ Since MP does not rely on any charge state determination, the masses of extensively glycosylated proteins can be readily measured. Advantages of MP include its rapid analysis time and a minimal need of sample preparation. A representative MP histogram of the SARS-CoV-2 S-trimer is depicted in **Figure 1A**. The S-trimer exhibits a large primary distribution at 474 kDa, while a minor low-mass distribution is also observed and can be assigned as residual S-monomer. Of note, no species corresponding to higher-order aggregates (i.e. dimers of S-trimers^25^) are observed.

**Figure 1.**
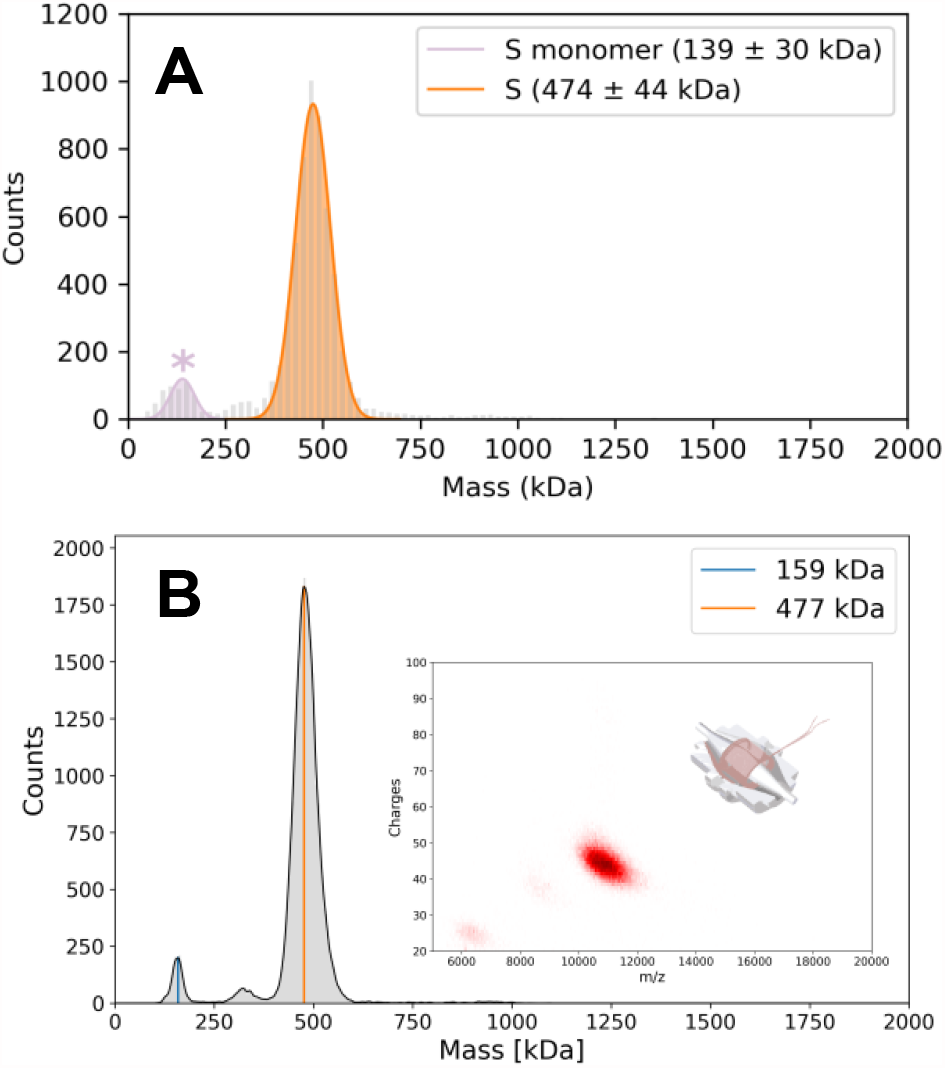
Representative mass histograms of the SARS-CoV-2 S-trimer. (A) MP histogram. (B) 1D CD-MS histogram, with the 2D CD-MS histogram shown in inset. The measured masses and abundances related to these data are provided in **Supplemental Table S1**.

Alternatively, charge detection mass spectrometry (CD-MS) can be used to overcome the charge inference problem in native MS by directly detecting both the charge and mass-to-charge (m/z) ratio of an ion.^42^ Due to this two-dimensional detection method, peaks that are unresolved in the m/z dimension may still be resolvable in the charge dimension, aiding in the assignment of complex spectra. A representative Orbitrap-based CD-MS histogram of the SARS-CoV-2 S-trimer is depicted in **Figure 1B**. Again, a single major distribution of particles corresponding to the S-trimer with is observed, with a minor distribution corresponding to the S-monomer also detected. The higher mass resolution achievable by CD-MS (as exhibited by the narrower mass distributions of the S-trimer relative to MP) highlight an important advantage of CD-MS. The trimer mass measured by CD-MS (477 kDa) is within ∼1% of the mass determined by MP. The close agreement in the results of these two disparate single particle methods underscores the robustness and complementarity of these approaches.

The backbone sequence-predicted mass of the S-trimer construct used here (390.349 kDa) underestimates the observed mass measured by both techniques by ∼90 kDa, reflecting the extensive glycosylation profile of the S protein. To estimate the expected mass contribution of the glycan shield, we calculated the average N-glycan masses derived from the glycoproteomic data of Allen and coworkers.^43^ The calculated glycan (92.0 kDa) and resultant total S-trimer (482.4 kDa) masses agree quite well (within 2%) with the masses measured by both MP and CD-MS. The glycan mass contribution measured here is somewhat lower than the recent results of Miller and co-workers^44^ who reported large mass discrepancies of ∼40% from similar glycoproteomic experiments. However, it should be noted the constructs used in that study differ from the one employed here in several key aspects (e.g. absence of stabilizing 2P mutations, different expression systems, etc.), as well as differing substantially in experimental setup (electrostatic linear ion trap vs. Orbitrap), which all may be factors accounting for this apparent discrepancy.

### Abs targeting the S-trimer can exhibit diverse binding characteristics

To establish the capability of single particle mass measurements to resolve the binding of Abs to the S-trimer, we initially screened the binding of a representative panel of twelve monoclonal anti-S-trimer IgGs using MP (**Figure 2**). These previously reported Abs, originally isolated from the sera of convalescent COVID-19 patients, target a variety of epitopes and exhibit varying neutralization potencies (**Supplemental Table S3**).^12^ Upon incubation of the S-trimer with the IgGs, new species of larger mass in the MP histograms are readily observed (**Figure 2A-B, Figure S1**). The evenly spaced, successive mass shifts of ∼150 kDa correspond to the binding of 1, 2, and 3 intact IgGs to the S-trimer. The particle distributions for each of the Abs is summarized as a heat map in **Figure 2C**.

**Figure 2.**
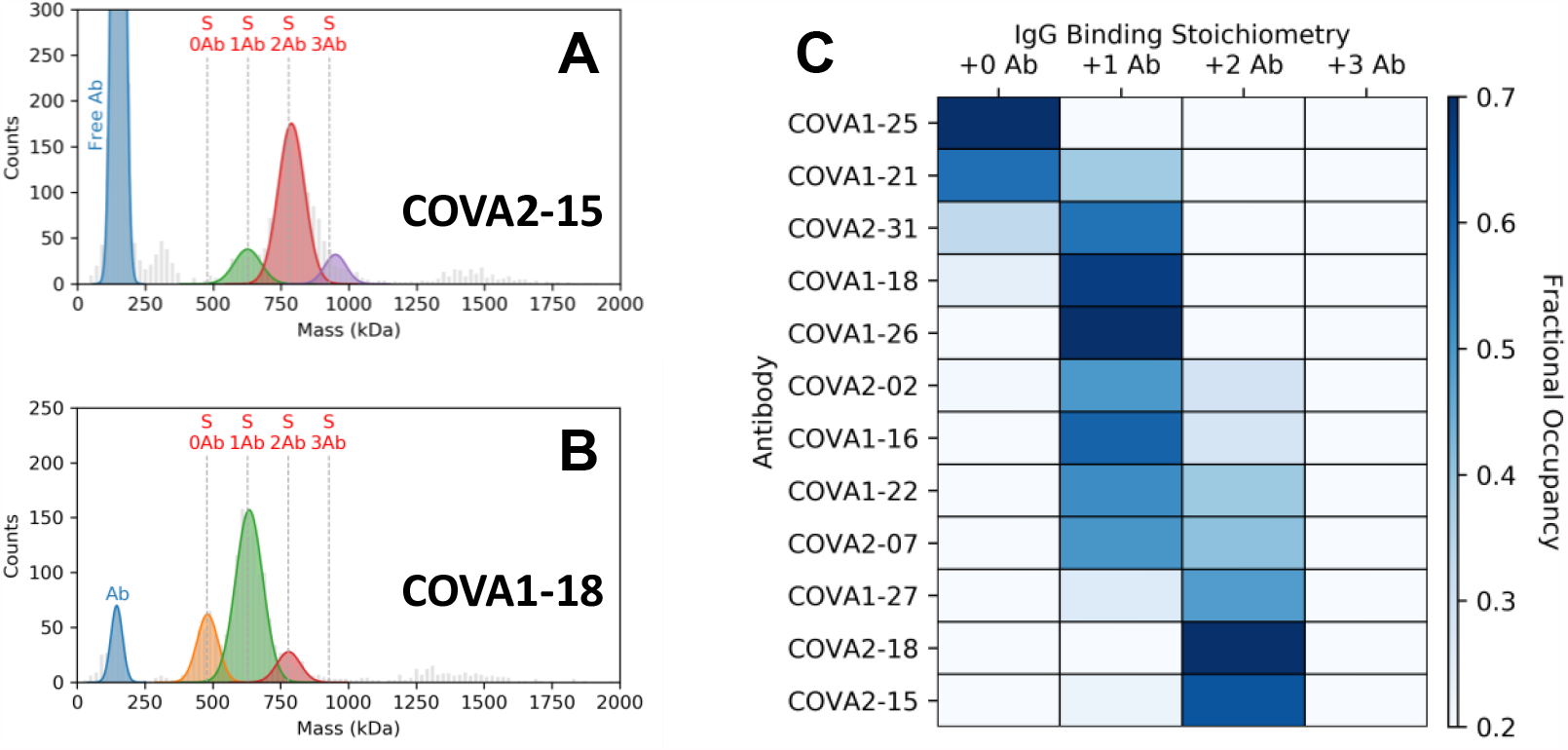
Measurement of IgG binding stoichiometries to the S-trimer by MP. MP histograms of the S-trimer following incubation with (A) COVA2-15 or (B) COVA1-18. The vertical dashed lines indicate the theoretical peak positions of each IgG-bound species. MP histograms of each of the Abs alone show a single major distribution at ∼150 kDa, in line with the expected IgG mass (**Figure S2Error! Reference source not found**.). The data clearly reveal that the “complete” 3:1 binding is not achieved for either Ab. COVA2-15 preferably binds two IgGs, whereas just one COVA1-18 binds to the S-trimer. Increasing concentrations of Ab do not change the preferred binding stoichiometries (**Figure S3**). Binding of both Abs to the S-trimer was also measured by CD-MS and very similar binding behaviour was observed, further illustrating the complementarity between MP and CD-MS (**Figure S4**). The low-abundance signals observed between 1200 and 1600 kDa originate from Ab-binding induced S-trimer dimers. (C) Fractional occupancies of each IgG-bound S-trimer species for a panel of twelve monoclonal Abs. A large diversity of binding stoichiometries are observed, ranging from 0 to 2. None of the tested Abs exhibited a preference for 3:1 binding. Additional representative MP histograms are depicted in **Figure S1**. A tabulation of binding stoichiometries related to these data are provided in **Supplemental Table S2**.

Our measurements reveal that Abs targeting the S-trimer can bind with a variety of preferred stoichiometries. Interestingly, none of the tested Abs exhibited a preference for the “complete” 3:1 (IgG:S-trimer) stoichiometry given the symmetry of the S-trimer. One may predict that these binding differences simply reflect different affinities of each Abs. Indeed, the two tested Abs with the lowest observed binding stoichiometries (COVA1-25 and COVA1-21) also have the weakest reported apparent dissociation constant (K_D,app_) values (>> 10 nM), and both exhibit a large proportion of free S-trimer. The remaining Abs, however, are quite similar in their affinities, with K_D,app_ values all in the sub-nM range (**Supplemental Table S3**). While COVA2-31, COVA1-18, COVA1-26, COVA2-02, COVA1-16, COVA1-22, and COVA2-07 preferably bound with a 1:1 stoichiometry, the dominant stoichiometry for COVA1-27, COVA2-18, and COVA2-15 was 2:1. The observation of diverse binding stoichiometries amongst the tested Abs, despite their very similar (and potent) K_D,app_ values, rules out affinity differences as the main driver of the remaining binding stoichiometries.

To help delineate other factors that may be modulating these stoichiometries, we next produced and evaluated Fab fragments and measured their binding to the S-trimer. Unlike the IgGs of each Ab, Fabs are only capable of binding one copy of an antigen (i.e. no avidity effects are possible), and due to their smaller size the contributions of steric clashes on the observed binding behaviour is minimal. These Fab experiments closely mimic previously reported analyses performed by single particle EM, where binding of Fab fragments was monitored.^11,25–27^ It is important to emphasize that while Fab fragments can clearly serve as a useful *in vitro* analogue, it is the intact IgG that is the biologically relevant species during the human immune response.

### COVA2-15 and COVA1-18

For these subsequent investigations, we focus specifically on two Abs: COVA2-15 and COVA1-18. These Abs, which both target epitopes on the RBD, were chosen firstly for their clinical relevance as both are among the most highly potent amongst the tested Abs in neutralizing the Wuhan SARS-CoV-2 strain, possessing near-identical neutralization potencies (IC_50_ ∼0.008 μg/mL).^12^ COVA1-18 has also been shown to protect cynomolgus macaques from high dose SARS-CoV-2 challenge.^10^ Secondly, despite these similar efficacies, our results indicate that these two Abs exhibit quite distinct (and representative) binding stoichiometries: COVA2-15 exhibits a preference for a 2:1 stoichiometry (with particles corresponding to 1, 2 or 3 bound IgGs, **Figure 2A**), whereas COVA1-18 displays a preference for 1:1 binding (with particles corresponding to 0, 1 or 2 bound IgGs, **Figure 2B**). In other words, the binding stoichiometries of these two Abs appear uncorrelated to both affinity and neutralization potency.

The binding behavior of the COVA1-18 and COVA2-15 Fabs differ substantially from that of their corresponding IgGs. When added in excess, the clearly observed mass shift reveals a preference for 3:1 binding for the COVA2-15 Fab by both MP (**Figure 3C**) and CD-MS (**Figure 3G-H**) – greater than the 2:1 seen for the full IgG. This stoichiometry agrees well with recent EM structures of the S-trimer bound to COVA2-15 Fabs, in which electron density for three bound Fabs was reported, and is in line with all three RBD copies of the S-trimer being occupied.^12^ Titration of COVA2-15 Fab at lower concentrations produce species of intermediate mass, corresponding to binding stoichiometries lower than 3:1 (**Figure 3A-B**). Interestingly, the COVA1-18 Fab exhibited essentially no binding to the S-trimer even when added in excess (**Figure 3F**), in contrast to the COVA1-18 IgG that revealed 1:1 binding (**Figure 2B**). This poor binding may explain why previous attempts to obtain a cryo-EM structure of COVA1-18 with the S-trimer using Fabs were unsuccessful (Andrew Ward, personal communication).

**Figure 3.**
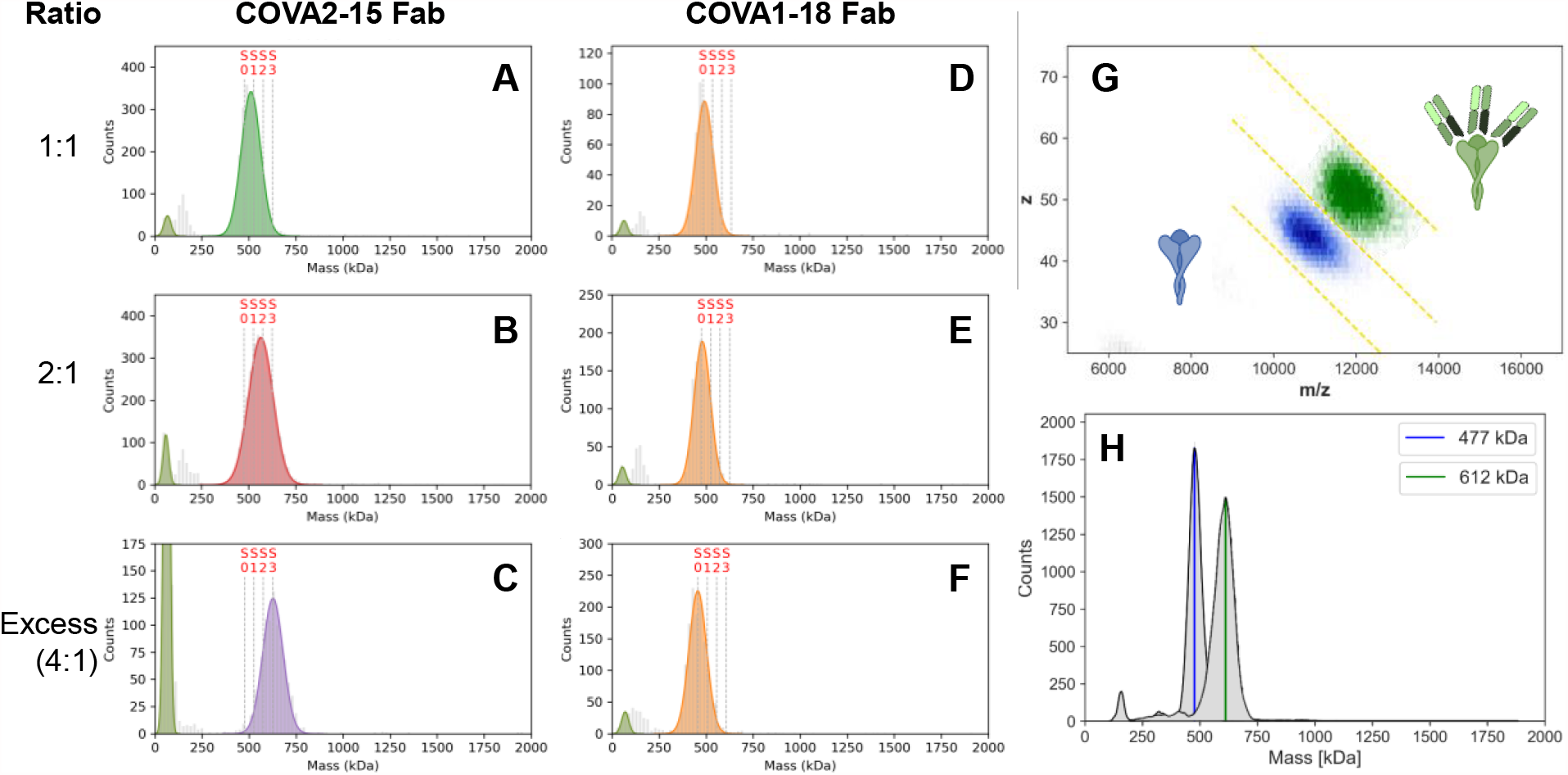
Stoichiometry of Fab binding to the S-trimer. (A-F) MP histograms of COVA2-15 and COVA1-18 Fab binding to the SARS-CoV-2 S-trimer at different mixing ratios. The vertical dashed lines indicate the theoretical peak positions of each Fab-bound stoichiometries. The data reveal that the S-trimer readily binds 3 COVA2-15 Fabs, whereas even in excess not a single COVA1-18 Fab binds to the S-trimer. (G-H) 2D and 1D CD-MS histograms of COVA2-15 Fab binding with excess ratio of Fab (green) as well as SARS-CoV-2 S-trimer only (blue). The observed shift in mass of ∼135 kDa confirms that the S-trimer predominantly binds 3 COVA2-15 Fabs.

What is the root cause of the divergent binding behaviors between both the different Abs, as well as between the IgGs and their associated Fabs? As described above, this could arise from several competing factors. For example, one may envision that the Ab binding stoichiometries could be simply reporting on the relative RBD up : down ratios in the S-trimer. Cryo-EM studies have suggested that the predominant states of the S-trimer are likely the [0 up : 3 down] and [1 up : 2 down] configurations.^11^ While this hypothesis has some qualitative agreement with the observed IgG binding stoichiometries (e.g. COVA1-18 would recognize the single “up” RBD state^12^, so a 1:1 stoichiometry is expected), it cannot satisfactorily rationalize either (1) the 2:1 binding seen in the COVA2-15 IgG (which binds agnostically to both “up” and “down” states^12^) nor (2) the different binding behaviour between the IgGs and Fabs. Evidently, other factors must also play a key role.

In the case of COVA1-18, the Fab displays substantially less binding than its corresponding IgG. This dramatic affinity loss going from intact IgG to Fab fragment is a hallmark of avidity (bivalent interactions).^21,26,45^ The possibility of avidity in the neutralization potency of COVA1-18 has recently been suggested, with measured K_D_ and pseudovirus IC_50_ values of the Fab more than 1 and 2 orders of magnitude worse, respectively, when compared to the full IgG.^10^ In the context of viral spike proteins, the bivalent IgGs can theoretically bind in two distinct modes: *inter*-spike (bridging between two different spike trimers) or *intra*-spike (binding two domains on the same spike).^21^ Distinguishing between different binding modes using standard biochemical assays that only monitor ensemble-averaged binding behaviour is not straightforward. By comparison, the mass measurements presented here readily allow differentiation of the two scenarios by their unique stoichiometries: *intra*-spike binding will produce Ab-bound species containing only one S-trimer, whereas *inter*-spike binding will produce species that will contain two S-trimers. Returning to **Figure 2**, the prominence of the [S + 1 Ab] species suggests that the *intra*-spike binding mode is the more prevalent mode for COVA1-18, although some signals in the 1200 to 1600 kDa range can be observed (which are absent in both the isolated S-trimer and in the presence of Fabs), suggesting a minor contribution of *inter*-spike binding is also possible. The lower-than-expected 1:1 binding stoichiometry seen in the COVA1-18 IgG then likely corresponds to a single Ab occupying two RBD binding sites on a single S-trimer due to bivalent binding (**Figure 4B**). Higher binding stoichiometries may then be inhibited due to the single available RBD site remaining (i.e. *intra*-spike binding is no longer possible).

**Figure 4.**
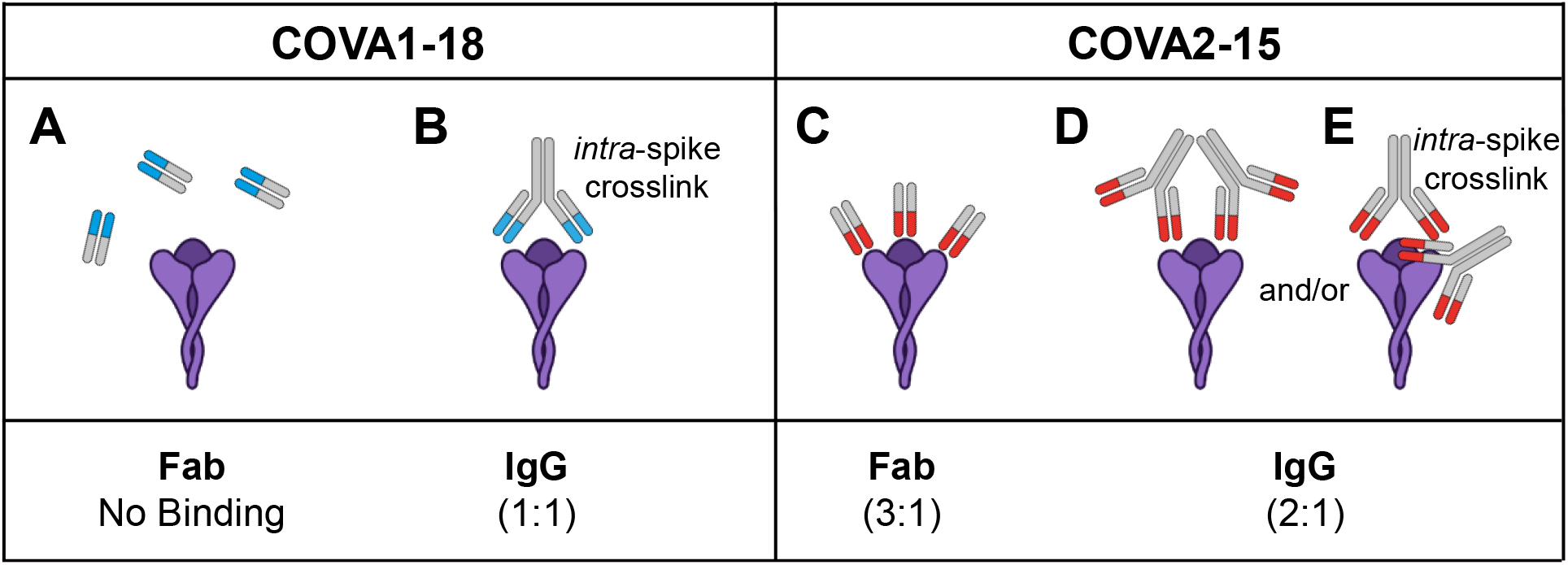
Proposed binding modes of COVA1-18 and COVA2-15 to the S-trimer. (A) For COVA1-18, its Fab has too low an affinity to effectively bind the S-trimer (violet). (B) In its native IgG format, bivalent interactions of the two Fabs enable effective binding with a dominant stoichiometry of 1:1. (C) For COVA2-15, its Fab possesses sufficient affinity alone to bind the S-trimer and occupies all three binding sites due to the lack of steric interactions. While the COVA2-15 IgG should theoretically be able to also bind with a 3:1 ratio, a combination of steric clashes (D) and/or bivalent binding (E) prevent this stoichiometry from being preferred.

For COVA2-15, the scenario is different as its Fab shows a higher binding stoichiometry than its corresponding IgG. One possibility is that binding of an initial IgG hampers the subsequent binding of additional IgGs (i.e. anti-cooperativity, **Figure 4D**). Considering that the smaller COVA2-15 Fab readily binds with the full 3:1 stoichiometry, the most likely source of this behaviour in this scenario would be steric clashes arising from the full IgG(s) that occlude the COVA2-15 IgG from fully occupying all three RBD sites. An alternative possibility is that COVA2-15, like COVA1-18, may also be capable of S-trimer binding *via intra*-spike crosslinking. In this scenario, one COVA2-15 IgG would bind bivalently to two RBD sites, while the remaining RBD site is occupied by a second, monovalently bound IgG (**Figure 4E**). This arrangement would also appear as a 2:1 binding stoichiometry, albeit with a different spatial configuration. Unlike COVA1-18, where avidity is a prerequisite for binding, in this arrangement COVA2-15 would seemingly not depend on this avidity to maintain affinity for the S-trimer, as evidenced by the binding capability of the COVA2-15 Fab (**Figure 3A-C**). Given that a small population of a 3:1 stoichiometry is observed for the COVA2-15 IgG (**Figure 2A**), it is likely that there exists a contribution of Fab-like, “monovalent-only” binding (**Figure 4D**) even if bivalent binding is the dominant binding mode (**Figure 4E**). Taken together, these results highlight the rich complexity inherent to IgG – S-trimer interactions, and the capacity of single particle analyses to aid in this unraveling this complexity.

### Sub-stoichiometric IgG binding is sufficient to prevent ACE2 binding

Given that COVA1-18 (and perhaps also COVA2-15) appears to leave at least one RBD site unoccupied, one may wonder if these Ab-bound S-trimers are still capable of binding the S-trimer host-receptor, ACE2. To explore this, we measured the binding of the ACE2 ectodomain against the S-trimer in the presence or absence of either the COVA1-18 or COVA2-15 IgG (**Figure 5**). ACE2-bound S-trimer species are distinguishable from their Ab-bound analogues by the different masses of the ACE2-dimer (200 kDa; **Figure 5A-C**) and an IgG (150 kDa; **Figure S2**). In the absence of any Ab, the S-trimer readily binds ACE2, with a predominant 1:1 stoichiometry at low mixing ratios as detected by MP (**Figure 5D**). MP measurements at higher ACE2 concentrations were partially impeded by spectral interference caused by a sub-population of a tetrameric ACE2 state which is of comparable mass to the free S-trimer (∼400 kDa vs. 477 kDa), although the species corresponding to ACE2-bound S-trimers remain unobstructed (**Figure S5**). While these species were also detected by CD-MS (and remain partially unresolved in the mass domain), the two species can be readily delineated in the 2D CD-MS histogram by their differences in both charge and m/z (**Figure 5E, H, and K**), highlighting the added potential of CD-MS to aid in interpreting spectrally congested data sets.

**Figure 5.**
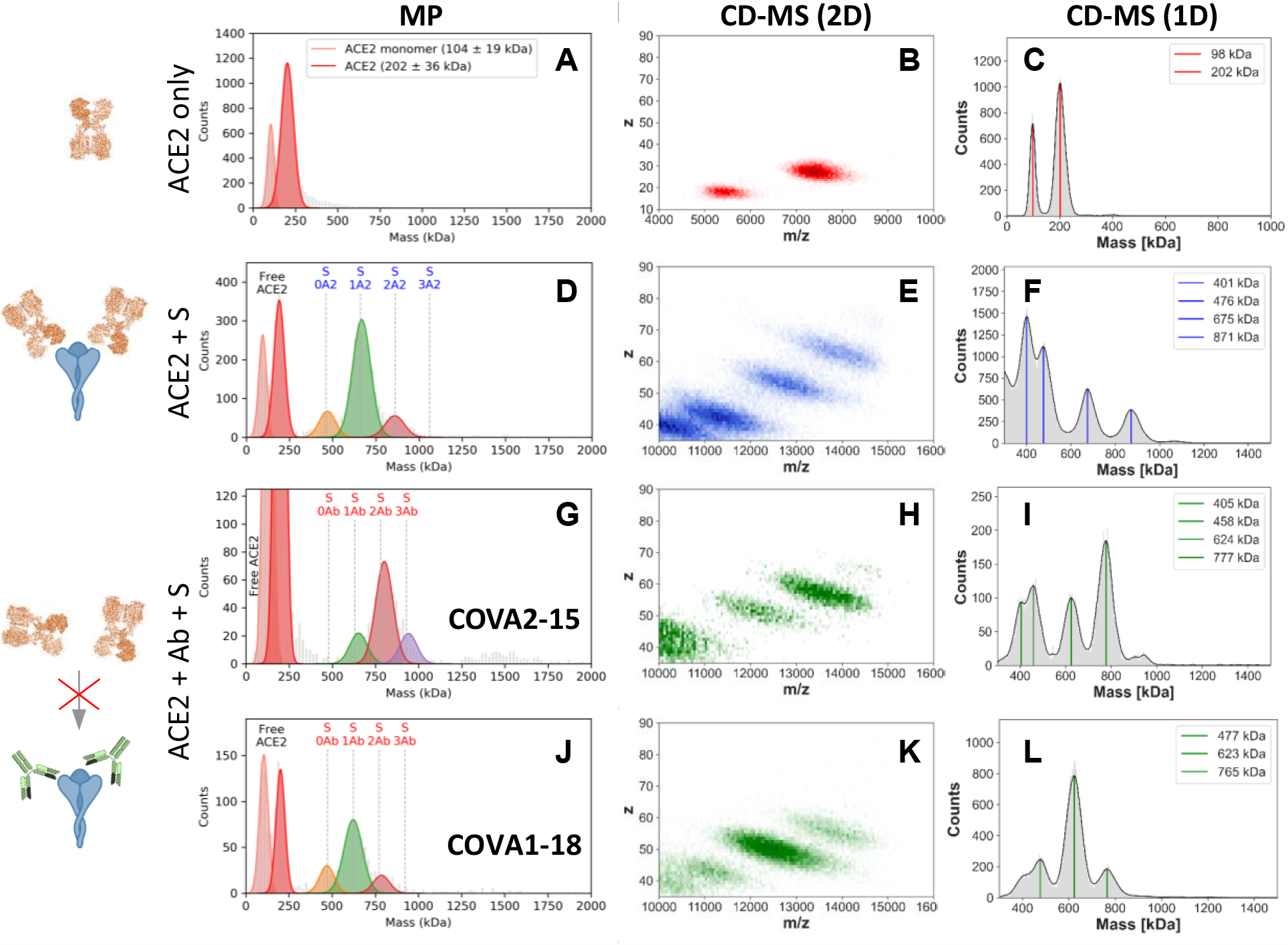
Sub-stoichiometric Ab binding to the S-trimer is sufficient to neutralize receptor binding. (A-C): MP and CD-MS histograms of ACE2 alone, revealing the dimeric nature of the utilized ACE2 construct and (D-F) ACE2 binding to the S-trimer. These results show that ACE2 is largely dimeric, and only the ACE2-dimer binds to S-trimer, whereby the S-trimer can accommodate either one or two ACE2. (G-L): MP and CDMS histograms of ACE2 binding to the S-trimer following pre-incubation with either COVA2-15 (G-I) or COVA1-18 (J-L). The observed mass shifts of ∼150 kDa (and not 200 kDa) indicate that both Abs fully prevent ACE2 binding to the S-trimer. Mixing ratios of 4:1 and 4:4:1 (ACE2-dimer : S-trimer and Ab : ACE2-dimer : S-trimer, respectively) were used for the CD-MS experiments, while 1:1 and 3:1:1 were used for the MP experiments. Note the similarities between the data presented in **Figure 2A/5G** and **2B/5J**.

In contrast to the clear observed binding of ACE2 in the absence of Ab, pre-incubation of the S-trimer with either the COVA2-15 or COVA1-18 IgG prior to addition of ACE2 produces only IgG-bound species, with no species observed corresponding to ACE2 binding, neither by formation of ternary [S-trimer + Ab + ACE2] complexes nor *via* displacement of bound Ab (**Figure 5G, J, I, and L**). It is likely that the same factors preventing the IgGs from reaching the “full” 3:1 stoichiometry (e.g. steric clashes and/or avidity effects) are preventing ACE2 from binding as well. Despite the seemingly available RBD site(s), it appears that sub-stoichiometric IgG binding is sufficient to fully block ACE2 binding, rendering them ideal neutralizing antibodies.

### Virus Variants of Concern

There is ongoing concern that newly emerging strains of the SARS-CoV-2 virus harboring additional mutations in the S protein may negatively impact the potency of already-existing anti-SARS-CoV-2 monoclonal Abs.^46–49^ As a proof-of-concept, we measured the binding of COVA2-15 and COVA1-18 against a S-trimer protein construct harboring the mutations present in the B.1.351 strain that originated in South Africa (**Figure 6**). In stark contrast to the original lineage, both Abs show substantially lower binding to this variant, with COVA1-18 exhibiting essentially no affinity. This binding loss is expected as COVA1-18 is unable to neutralize B.1.351, while COVA2-15 has substantially reduced activity.^10,50^ These results strengthen the arguments for the necessity of using multiple Abs (cocktails) for the design of target biotherapeutic treatments, and also highlight the potential of mass photometry and charge detection mass spectrometry to guide Ab design and development.

**Figure 6.**
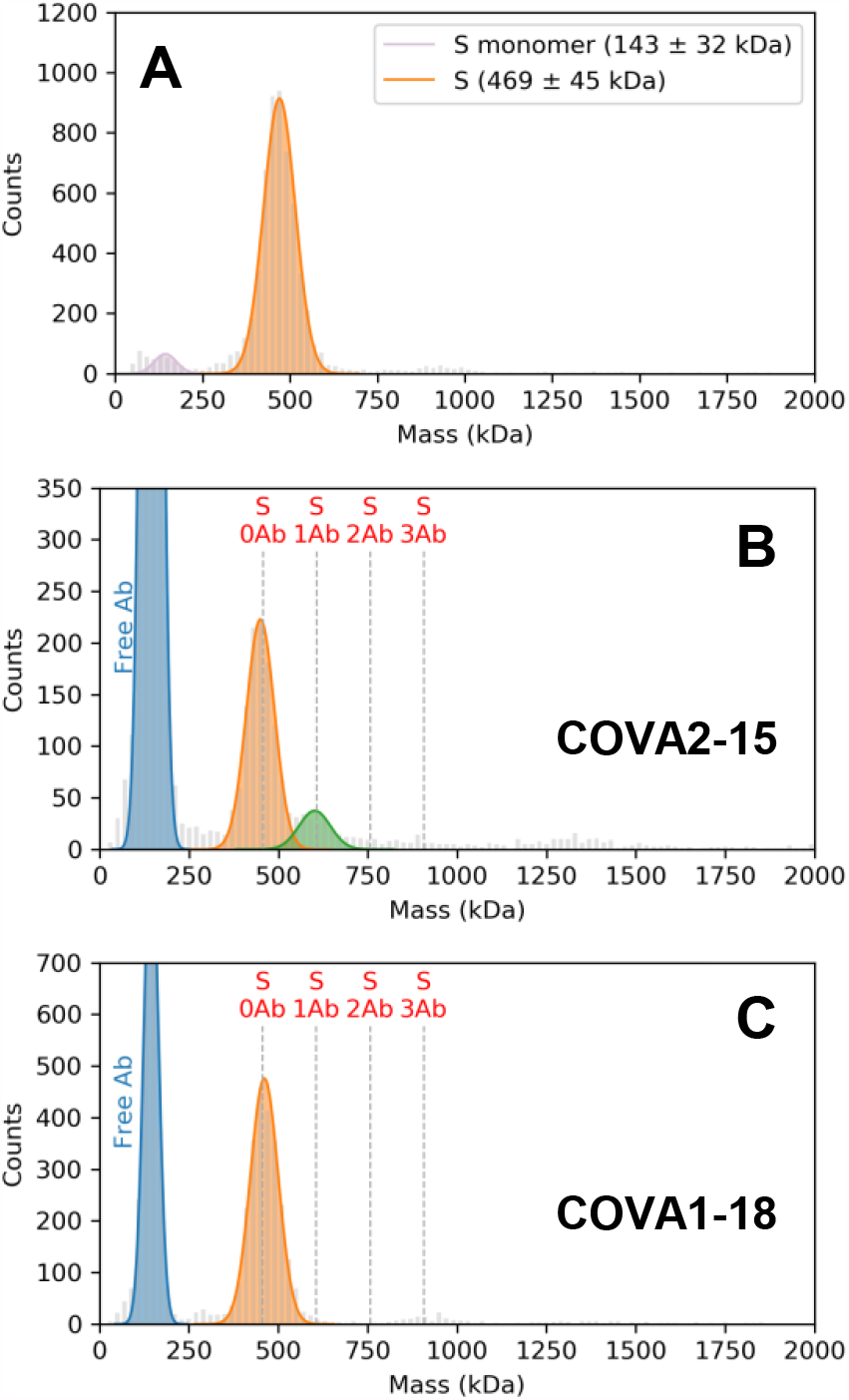
MP histograms of COVA2-15 and COVA1-18 binding to the SARS-CoV-2 variant N501Y.V2 S-trimer. (A) Variant S-trimer alone. S-trimer incubated with COVA2-15 (B) or COVA1-18 (C). In stark contrast to the original lineage (**Figure 2**), essentially no Ab binding is observed.

## Discussion

We demonstrate here the unique application of two single particle approaches, MP and CD-MS, for interrogating the interaction stoichiometries between full Abs, the ACE2 receptor, and the SARS-CoV-2 S protein ectodomain. We find that different Abs can exhibit surprisingly distinct binding behavior. In the case of the potent neutralizing Abs COVA2-15 and COVA1-18, different binding stoichiometries can arise despite commonly targeting the RBD and having identical neutralization potencies. This behaviour is not fully recapitulated when analyzing the binding of Fab fragments, stressing the necessity of studying Ab-antigen interactions in the context of the full, non-truncated IgG. Our results highlight the complex interplay of affinity, avidity, and anti-cooperativity effects in these interactions, and the capability of single particle mass analysis to shed light on these co-occurring phenomena.

Our analyses here focus primarily on the binding behaviour of the two representative neutralizing Abs COVA2-15 and COVA1-18. One may wonder if the determinants of the 1:1 and 2:1 binding behaviour that we uncovered for these Abs can be generalized to other anti-S-trimer Abs (e.g. **Figure 2C**). While it is tempting to speculate, for example, that all 1:1 binding IgGs bind in a manner analogous to COVA1-18 (i.e. bivalently), in reality the situation may be more complex. Other factors such as steric blockage, incompatible angles of approach, or the location of the epitope cannot be dismissed *a priori*. As such, the binding determinants of each Ab should be determined on a case-by-case basis. Nevertheless, the experimental approaches outlined in this work, especially in combination with already-established methods such as single particle EM, are well-suited to address these questions.

Our investigations were enabled by the capacity of recently developed single particle approaches to overcome the high degree of mass spectral complexity normally brought by the extensive glycosylation of the SARS-CoV-2 S protein. We expect that these technologies will open the door for studies into similarly complex biological systems, such as glycoproteins from other viruses and biological agents. We foresee that these techniques will be especially useful in the characterization and rational design of biotherapeutics, e.g. monoclonal Ab cocktails or multivalent nanobodies.^51,52^ It is anticipated that single particle mass analysis will provide a powerful addition to the toolbox of contemporary biophysical methods to study protein-protein interactions.

## Acknowledgements

This research received funding through the Dutch Research Council (NWO) funding the Netherlands Proteomics Centre through the X-omics Road Map program (project 184.034.019). AJRH acknowledges support from the Netherlands Organization for Scientific Research (NWO) through the Spinoza Award SPI.2017.028 to AJRH. RWS acknowledges support from the Netherlands Organization for Scientific Research (NWO) through a Vici grant. The authors also acknowledge support from the Bill & Melinda Gates Foundation grants INV-002022 and INV-008818 (to RWS), INV-024617 (MJvG) and INV-004923 (to IAW). MJvG is a recipient of an Amsterdam UMC AMC Fellowship.

## Author Contributions

Project Conception: VY, SHL, RWS, MJvG, AJRH

Protein Synthesis: TCG, PJMB, MB, YA, HL, MY

Data Collection: VY, SHL

Writing: VY, SHL, RWS, MJvG, IAW, AJRH

Supervision: IAW, RWS, MJvG, AJRH

Editing / Review: All authors

## Materials and Methods

### WT and B.1.351 spike proteins, human ACE2 receptor, and antibodies

The 2P-stabilized S proteins of the Wuhan strain (WT) and B.1.351 variant were described previously.^12,50^ The B.1.351 construct contained the following mutations compared to the WT variant (Wuhan Hu-1; GenBank: MN908947.3): L18F, D80A, D215G, L242H, R246I, K417N, E484K, N501Y, D614G and A701V. Both S constructs were produced in HEK293F suspension cells (ThermoFisher) and purified as previously described.^12^ For the human ACE2 receptor, soluble ACE2 was generated as described previously^12^ by using a gene encoding amino acids 18-740 of ACE2. The IgGs and Fab fragments used in this study were produced as previously described.^12,26^

### Mass Photometry

MP experiments were performed on a Refeyn OneMP (Refeyn Ltd.). 24 mm x 50 mm microscope coverslips (Paul Marienfeld GmbH) were cleaned by serial rinsing with Milli-Q water and HPLC-grade isopropanol (Fisher Scientific Ltd.), on which a CultureWell gasket (Grace Bio-labs) was then placed. For each measurement, 12 µL of buffer was placed in the well for focusing, after which 3 µL of sample was introduced and mixed. Movies were recorded for 120 seconds at 100 fps under standard settings. MP measurements were calibrated using an in-house prepared protein standard mixture: IgG4Δhinge-L368A (73 kDa^53^), IgG1-Campath (149 kDa), Apoferritin (479 kDa), and GroEL (800 kDa). MP data were processed using DiscoverMP (Refeyn Ltd).

All MP measurements were performed in Tris buffer (25 mM Tris, 100 mM NaCl, pH 7.6 (Sigma Aldrich)). For each experiment, a 100 nM solution of SARS-CoV-2 S protein was mixed with an equal volume of ligand to the desired concentration ratio and incubated at room temperature (22 °C) for 5 minutes. Unless otherwise stated in the text, ligands were mixed at a 3:1 (ligand : S-trimer) molar ratio. Afterwards, 3 µL of the reaction mixture was immediately transferred to the instrument for measurement. For binding experiments containing both Abs and ACE2, S protein was pre-incubated with Ab for 5 minutes as described above, after which an equal volume of ACE2 solution at the desired concentration was added and incubated for a further 5 minutes prior to loading onto the instrument.

### CD-MS

CD-MS measurements were performed on an Orbitrap Q Exactive UHMR mass spectrometer (Thermo Fisher Scientific). Samples were introduced into a gold-coated borosilicate capillary (prepared in-house) for nano-electrospray ionization in positive ion mode. A resolution of 200,000 at 400 m/z was set for 1 s ion transient. The noise level parameter was fixed at 0. Nitrogen was used as collision gas. The in-source-trapping voltage and HCD voltage were optimized for maximal ion transmission. After multi scan acquisition, .RAW files were centroided and converted into mzXML format for processing as previously described.^39^ A calibration factor of 12.55 (normalized arbitrary intensities/charges) was applied for correlating the measured intensities and charges of individual single ions. Several mzXML files could be merged to one for providing larger number of statistics. According to the determined charge state, a resulting formula Mass = m/z * z -z was used to calculate the mass of each single ion respectively.

All samples for CD-MS measurements were first buffer exchanged into 500 mM ammonium acetate solution (pH 7.5) using Amicon 10 kDa MWCO centrifugal filters (Merck Millipore), unless otherwise stated. For IgG binding experiments, a 100 nM solution of SARS-CoV-2 S protein was mixed with an equal volume of ligand to an excess ratio (4 Abs : 1 S-trimer) and incubated at room temperature (22 °C) for at least 5 minutes. Afterwards, ∼3 µL of the reaction mixture was introduced into the mass spectrometer for the measurement. For binding experiments containing both Abs and ACE2, S protein was pre-incubated with Ab for 5 minutes as described above, after which an equal volume of ACE2 solution at the desired concentration was added and incubated for a further 5 minutes prior to loading onto the instrument. For Fab binding experiments, a 1 µM solution of SARS-CoV-2 S protein was mixed and pre-incubated with an equal volume of Fab to an excess ratio (5 Fabs : 1 S-trimer) before buffer exchange into 500 mM ammonium acetate solution (pH 7.5) using a Micro Bio-Spin 6 column (Bio-Rad).

## Supplemental Figures

**Figure S1.**
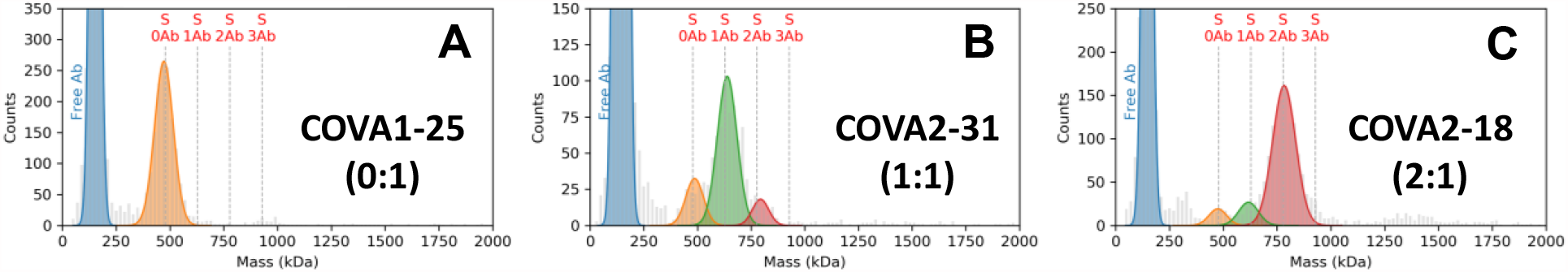
Representative MP histograms of three tested Abs (IgG format) bound to the S-trimer. (A) COVA1-25 (0:1 stoichiometry). (B) COVA2-31 (1:1 stoichiometry). (C) COVA2-18 (2:1 stoichiometry). As demonstrated here, different anti-S-trimer Abs can exhibit a variety of binding stoichiometries.

**Figure S2.**
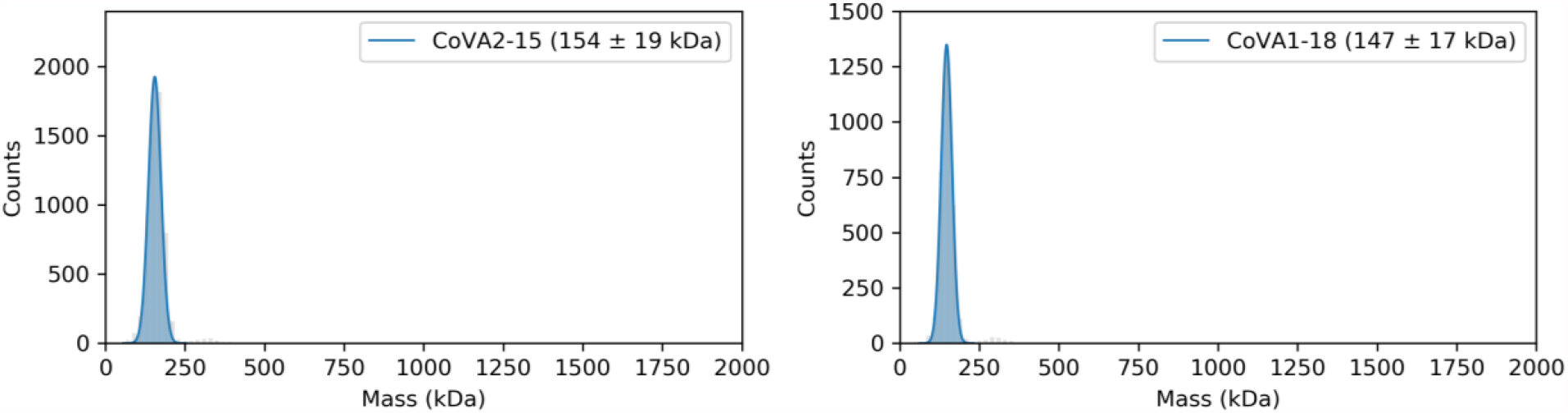
MP histograms of the COVA2-15 and COVA1-18 IgGs. A single mass distribution at ∼150 kDa is observed for both Abs in line with the predicted masses.

**Figure S3.**
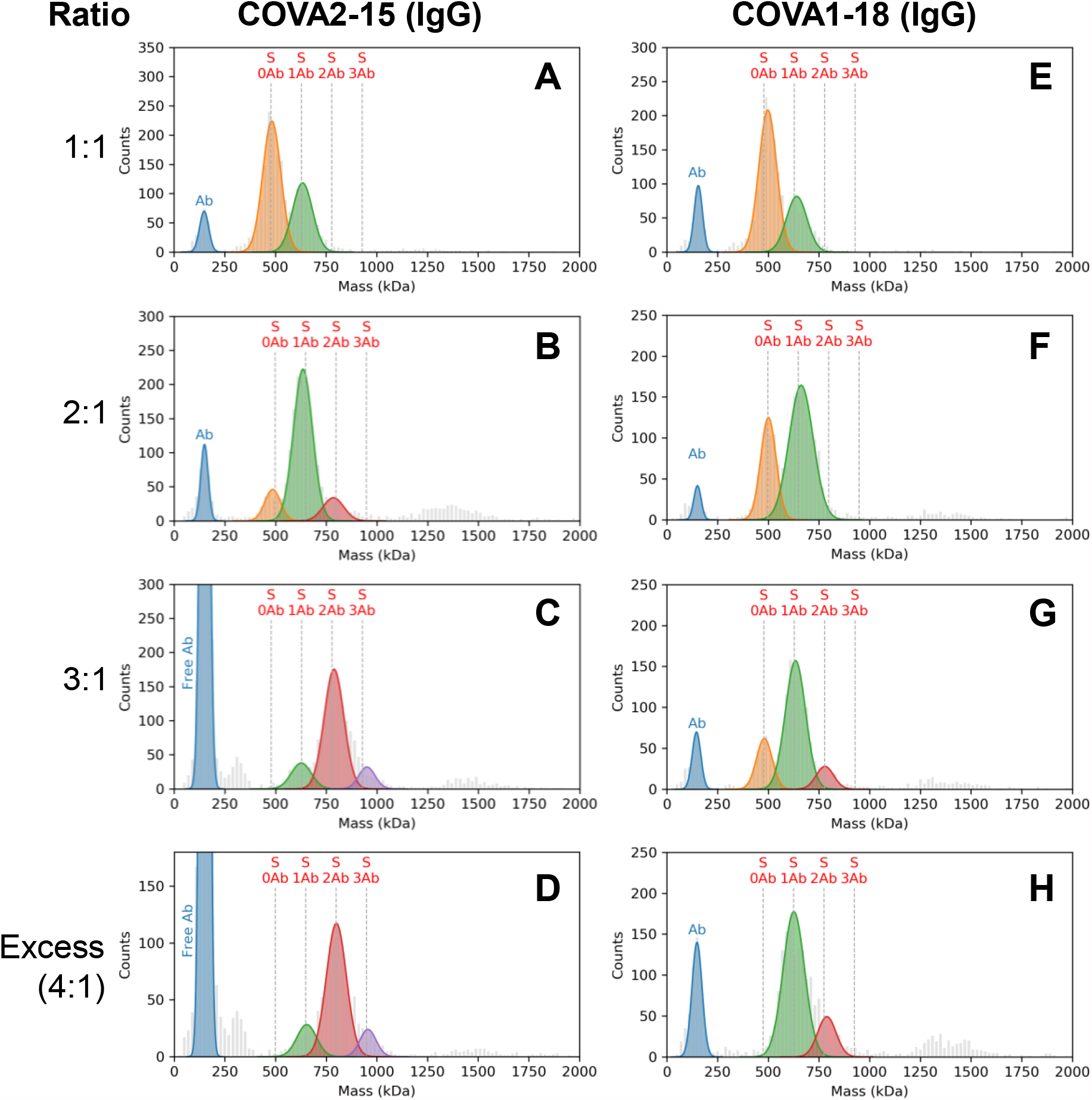
Stoichiometry of COVA2-15 and COVA1-18 binding to the S-trimer assessed by MP. MP histograms of COVA2-15 and COVA1-18 IgG binding to the SARS-CoV-2 S-trimer at different mixing ratios. (A-D): COVA2-15 Ab. (E-H): COVA1-18 Ab. The vertical dashed lines indicate the theoretical peak positions of each Ab-bound species. The data clearly reveal that full stoichiometric binding is not achieved for either Ab, but also that COVA2-15 preferably binds two Abs, whereas even at excess preferably just one COVA1-18 binds to the S-trimer. As expected, lower mixing ratios result in lower observed binding stoichiometries. The low-abundance signals observed between 1200 and 1600 kDa originate from Ab-binding induced S-trimer dimers. The measured masses and abundances related to these data are provided in **Supplemental Table S4**.

**Figure S4.**
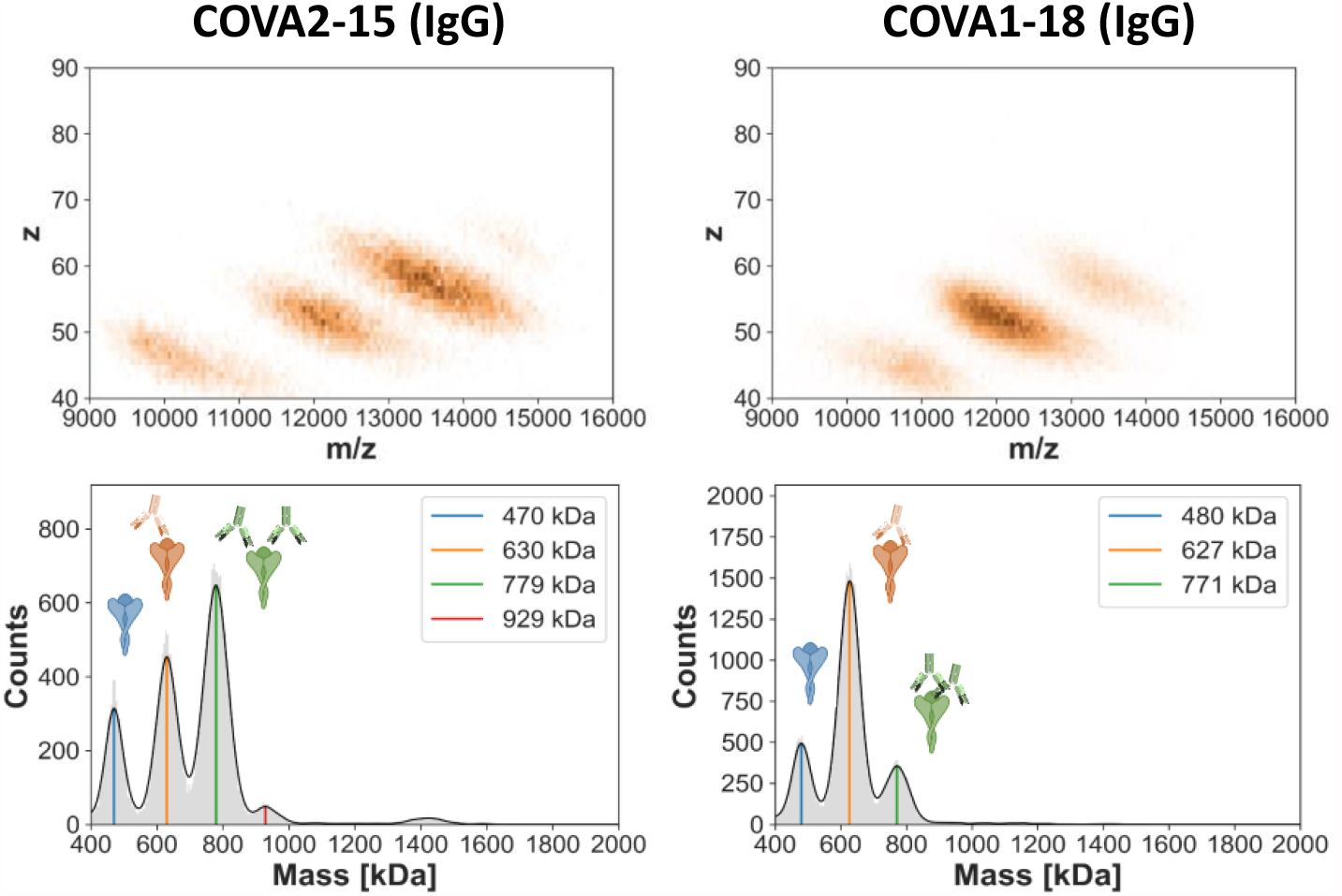
Stoichiometry of Ab binding to the S-trimer assessed by Orbitrap-based charge detection native mass spectrometry. Two-dimensional mass spectra of COVA2-15 and COVA1-18 Ab binding to the SARS-CoV-2 S-trimer at a mixing ratio of 4:1. Left: COVA2-15 Ab. Right: COVA1-18 Ab. The data are in line with the MP data presented in **Figure 2**, and also reveal that full stoichiometric binding is not achieved for either Ab, but also that COVA2-15 binds more readily two Abs, whereas even at access preferably just one COVA1-18 binds to the S-trimer. The measured masses and abundances related to these data and provided in **Supplemental Table S4**.

**Figure S5.**
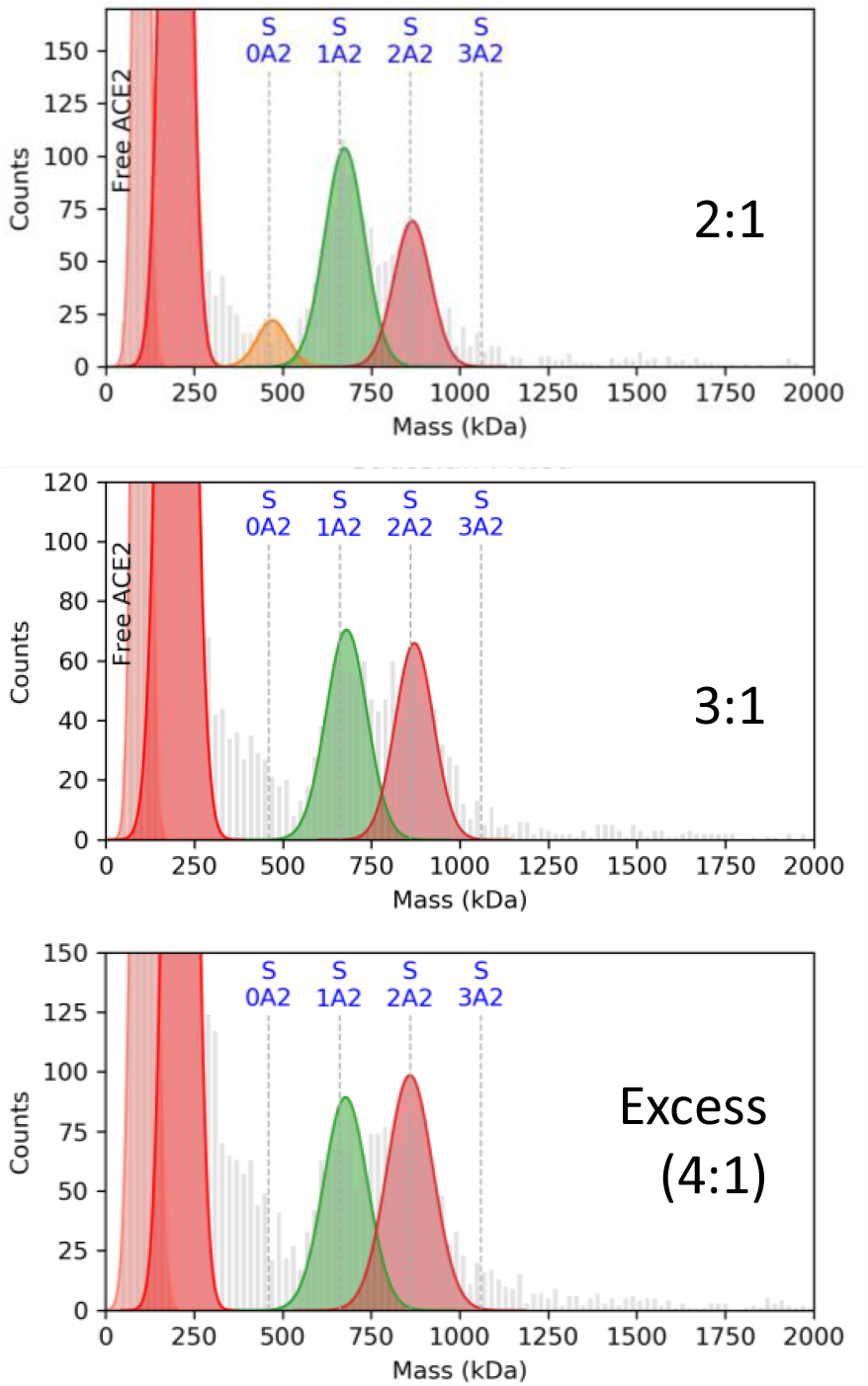
MP histograms of ACE2 binding to the S-trimer at increasing ratios. Even at an excess of 4:1, a substantial number of S-trimers still only bind one ACE2, and essentially no binding of 3 ACE2 is observed. Interference from a tetrameric ACE2 sub-population occlude accurate quantitation of the free S-trimer.

**Supplemental Table S1.** Tabulation of masses of SARS-CoV-2 S-trimer.

| Method | Species | Mass (kDa) | FWHM (kDa) | Normalized Abundance (%) |
| --- | --- | --- | --- | --- |
| MP | S (monomer) | 139 | 73 | 8 |
| MP | S (trimer) | 475 | 106 | 92 |
| CD-MS | S (monomer) | 159 | 29 | 5 |
| CD-MS | S (trimer) | 477 | 61 | 95 |

**Supplemental Table S2.** Tabulation of fractional occupancies of each Ab bound to the S-trimer. Tabulated values are derived from the normalized summation of two 120 second MP acquisitions.

| IgG | Fractional Occupancies |  |  |  |
| --- | --- | --- | --- | --- |
|  | +0 Ab | +1 Ab | +2 Ab | +3 Ab |
| COVA1-25 | 1.000 | 0.000 | 0.000 | 0.000 |
| COVA1-21 | 0.579 | 0.384 | 0.037 | 0.000 |
| COVA2-31 | 0.334 | 0.572 | 0.095 | 0.000 |
| COVA1-18 | 0.249 | 0.671 | 0.080 | 0.000 |
| COVA1-26 | 0.127 | 0.710 | 0.164 | 0.000 |
| COVA2-02 | 0.211 | 0.497 | 0.292 | 0.000 |
| COVA1-16 | 0.112 | 0.601 | 0.287 | 0.000 |
| COVA1-22 | 0.090 | 0.522 | 0.388 | 0.000 |
| COVA2-07 | 0.089 | 0.503 | 0.408 | 0.000 |
| COVA1-27 | 0.182 | 0.266 | 0.491 | 0.062 |
| COVA2-18 | 0.115 | 0.127 | 0.711 | 0.047 |
| COVA2-15 | 0.023 | 0.228 | 0.628 | 0.121 |

**Supplemental Table S3.** Tabulation of Ab epitopes, neutralization, and K_D,app_.

| **Antibody** | **Epitope\*** | **Neutralizes?\*** | **$K_{D,app}$**<br>**(nM)\*** |
| --- | --- | --- | --- |
| COVA1-25 | non-RBD | Yes | N.D. |
| COVA1-21 | non-RBD | Yes | 34 |
| COVA2-31 | RBD | No | 0.3 |
| COVA1-18 | RBD | Yes | 0.03 |
| COVA1-26 | NTD | No | 0.9 |
| COVA2-02 | RBD | Yes | 0.3 |
| COVA1-16 | RBD (Up) | Yes | 0.3 |
| COVA1-22 | NTD | Yes | 0.4 |
| COVA2-07 | RBD | Yes | 0.6 |
| COVA1-27 | non-RBD | No | 0.7 |
| COVA2-18 | S2 | No | 1.6 |
| COVA2-15 | RBD (Up+Down) | Yes | 0.6 |

\* Data from ref<sup>12</sup>

**Supplemental Table S4.**
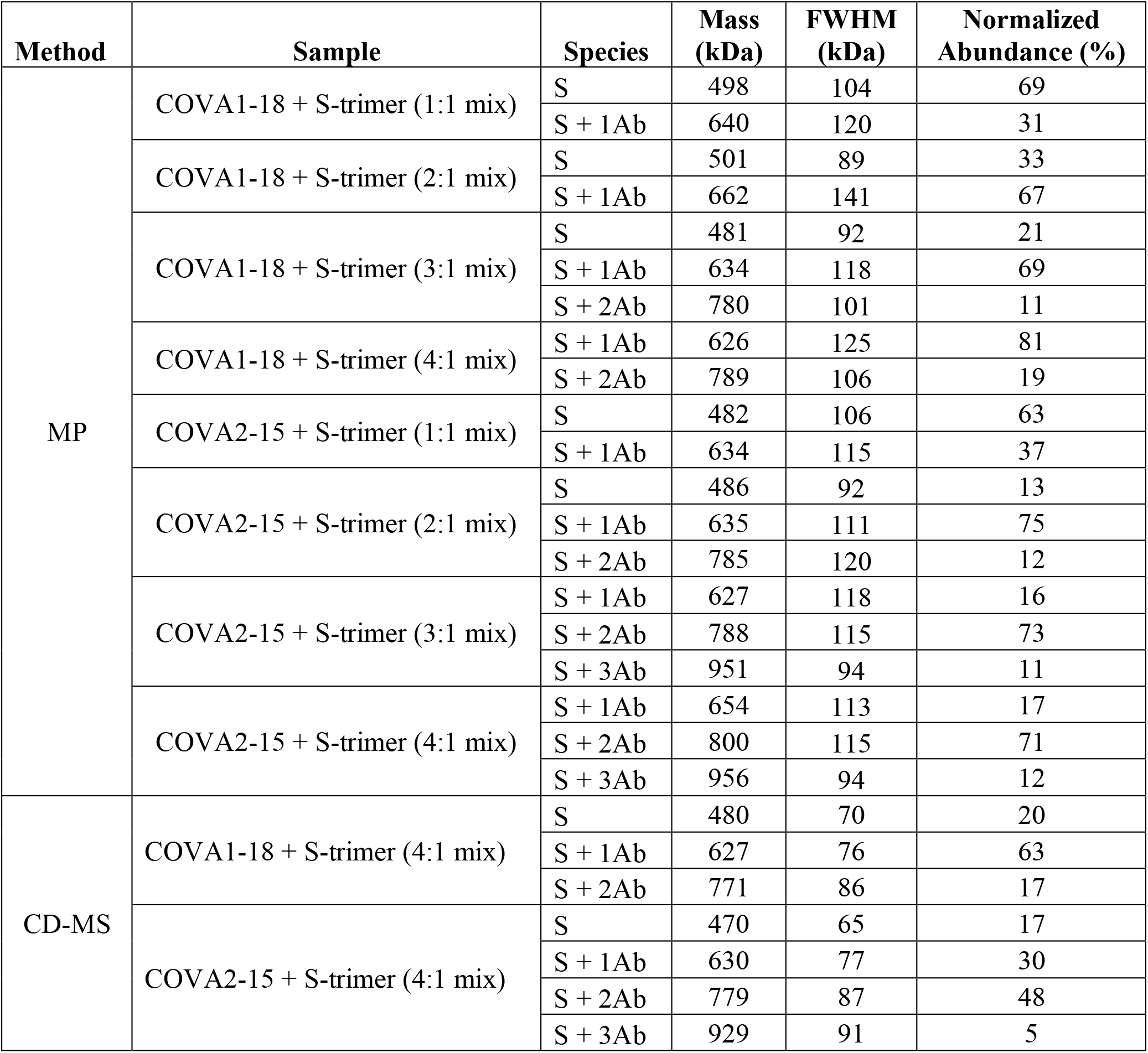
Tabulation of masses for Ab binding experiments.

| Method | Sample | Species | Mass (kDa) | FWHM (kDa) | Normalized Abundance (%) |
| --- | --- | --- | --- | --- | --- |
| MP | COVA1-18 + S-trimer (1:1 mix) | S | 498 | 104 | 69 |
|  |  | S + 1Ab | 640 | 120 | 31 |
|  | COVA1-18 + S-trimer (2:1 mix) | S | 501 | 89 | 33 |
|  |  | S + 1Ab | 662 | 141 | 67 |
|  | COVA1-18 + S-trimer (3:1 mix) | S | 481 | 92 | 21 |
|  |  | S + 1Ab | 634 | 118 | 69 |
|  |  | S + 2Ab | 780 | 101 | 11 |
|  | COVA1-18 + S-trimer (4:1 mix) | S + 1Ab | 626 | 125 | 81 |
|  |  | S + 2Ab | 789 | 106 | 19 |
|  | COVA2-15 + S-trimer (1:1 mix) | S | 482 | 106 | 63 |
|  |  | S + 1Ab | 634 | 115 | 37 |
|  | COVA2-15 + S-trimer (2:1 mix) | S | 486 | 92 | 13 |
|  |  | S + 1Ab | 635 | 111 | 75 |
|  |  | S + 2Ab | 785 | 120 | 12 |
|  | COVA2-15 + S-trimer (3:1 mix) | S + 1Ab | 627 | 118 | 16 |
|  |  | S + 2Ab | 788 | 115 | 73 |
|  |  | S + 3Ab | 951 | 94 | 11 |
|  | COVA2-15 + S-trimer (4:1 mix) | S + 1Ab | 654 | 113 | 17 |
|  |  | S + 2Ab | 800 | 115 | 71 |
|  |  | S + 3Ab | 956 | 94 | 12 |
| CD-MS | COVA1-18 + S-trimer (4:1 mix) | S | 480 | 70 | 20 |
|  |  | S + 1Ab | 627 | 76 | 63 |
|  |  | S + 2Ab | 771 | 86 | 17 |
|  | COVA2-15 + S-trimer (4:1 mix) | S | 470 | 65 | 17 |
|  |  | S + 1Ab | 630 | 77 | 30 |
|  |  | S + 2Ab | 779 | 87 | 48 |
|  |  | S + 3Ab | 929 | 91 | 5 |

